# Characterisation of naturally occurring MERS-CoV Spike mutations and their impact on entry and neutralisation

**DOI:** 10.64898/2026.02.17.706312

**Authors:** Rachael Dempsey, Hannah Goldswain, Joseph Newman, Nazia Thakur, Tracy MacGill, Todd Myers, Robert Orr, Dalan Bailey, James P. Stewart, Waleed Aljabr, Julian A. Hiscox

## Abstract

In this study the phenotypic consequences of naturally occurring single nucleotide polymorphisms (SNPs) in the MERS-CoV Spike protein were investigated. The impact of Spike mutations on virus entry and neutralisation of contemporary MERS-CoV strains is not currently well understood. Naturally occurring mutations were identified by aligning 584 MERS-CoV Spike sequences from either human clinical isolates collected between 2012 – 2024 or from viruses passaged in human cells. Fifteen SNPs of interest occurring in the NTD, RBD and adjacent to the S1/S2 cleavage site were selected for further characterisation based on their location in the Spike protein, frequency and identification in previous studies. A representative clade B, lineage 5 wildtype Spike sequence, which reflected those carried by MERS-CoV viruses circulating in the Middle East, was used in this study. The mutations of interest were introduced to the wildtype backbone to generate Spike variants. A lentiviral-based pseudotyping system was then used to investigate the impact of these Spike mutations on entry and neutralisation. I529T, E536K and L745F were shown to improve MERS-CoV entry. L411F, T424I, L506F, L745F and T746K were found to increase resistance to neutralisation by pooled patient sera. This study has identified novel naturally occurring Spike mutations that resulted in phenotypic differences in virus entry and neutralisation of contemporary MERS-CoV strains. Continued investigation of the phenotypic consequences of naturally occurring MERS-CoV Spike mutations is essential for assessing the risk to public health, especially given the pandemic potential of this virus.

**Importance:** The main aim of this study was to investigate the impact of naturally occurring MERS-CoV Spike mutations on virus entry and neutralisation. The phenotypic consequences of mutations occurring in the Spike protein of contemporary MERS-CoV strains are not currently well understood. Improving our understanding is of particular importance due to MERS-CoV continuing to pose a public health risk, with frequent spillover events and mounting evidence of human-to-human transmission since the virus emerged in 2012. A major concern is that as MERS-CoV continues to evolve, it may become more infectious, resulting in increased transmission between humans. To add to this, surveillance is limited and there are currently no specific medical countermeasures available to treat MERS-CoV disease. The MERS-CoV Spike pseudotyping system developed in this study is a useful tool that could be used alongside surveillance systems to rapidly assess novel Spike mutations in functional assays. This MERS-CoV pseudotyping system could also be used to aid the development of medical countermeasures such as vaccines, antivirals and antibody therapies.

## Introduction

MERS-CoV evolves via recombination, single nucleotide polymorphisms (SNPs), insertions and deletions (indels) (1). Since emerging in 2012, three genetically distinct clades have been identified; clades A, B and C (2–4). The MERS-CoV Erasmus Medical Centre/2012 (EMC/2012) reference strain, isolated from the first patient identified as being infected with the virus in 2012, was phylogenetically classified as clade A and has been the focus of most MERS-CoV research studies to date. However, clade A viruses were outcompeted by clade B viruses and have not been detected since 2015 (4, 5). Clade B viruses continue to circulate in dromedary camels and humans, predominantly in the Middle East (4, 6), and have been the cause of epidemic outbreaks of MERS-CoV since 2012 (7). The most notable outbreaks in humans occurred in Saudi Arabia in 2014 and South Korea in 2015 (8, 9), where the index case was a traveller returning from the Middle East (9–11). Human-to-human transmission was observed during these outbreaks (9–12), which were associated with a recombination event between clade B viruses belonging to lineages 3 and 4 (13). The recombination event occurred at ORF1ab/Spike and resulted in a new lineage: clade B lineage 5. Lineage 5 viruses were of superior replicative fitness and caused lower levels of cytokine and interferon (IFN) induction relative to lineages 3 and 4, which may have resulted in the increased human cases during these outbreaks (7, 13). Clade C viruses are predominantly present in camels in African countries, with minimal human infections being observed for this clade (2, 3, 14).

Clade B viruses have been phylogenetically classified into seven lineages, indicating genome diversity and continued evolution of the viruses in this clade (4). SNPs occurring in Spike are of particular interest due to this protein mediating MERS-CoV entry into host cells. Spike is also the main target of neutralising antibodies (15) and SNPs in the gene that encodes for this protein may lead to partial immune escape in individuals that have previously experienced a natural MERS-CoV infection. Additionally, Spike is a target for both human and camel vaccines being trialled to protect against MERS-CoV infection (16–21). Therefore, changes in Spike may confer an advantage to the virus by improving entry and making MERS-CoV more infectious (22), or by leading to partial immune escape and reducing immunity in individuals previously infected with MERS-CoV (23). Improved entry and/or partial immune escape of MERS-CoV could result in increased virus transmission in humans, which is a major public health concern.

The Spike protein exists as a trimer at the surface of the viral envelope, with each monomer carrying a non-covalently associated S1 and S2 subunit after cleavage at S1/S2 (24–26). The S1 subunit contains the N-terminal domain (NTD) responsible for attachment to the host cell (27–29) and the receptor binding domain (RBD) that interacts with Dipeptidyl peptidase-4 (DPP4) receptor (30, 31). The S2 subunit mediates membrane fusion and contains the S2’ cleavage site, fusion peptide, heptad repeat domains 1 and 2, transmembrane domain and cytoplasmic domain (26). Spike binds the DPP4 receptor on the host cell surface via its S1 subunit, which subsequently leads to cleavage at S2’ (26, 30). This cleavage event enables the fusion of viral and host membranes via the S2 subunit and entry of the virus into the host cell (26). Entry into the host cell can be prevented by neutralising antibodies binding to the virus. The majority of MERS-CoV neutralising antibodies bind to the S1 subunit, usually to the RBD, although NTD-specific neutralising antibodies have also been identified (15). Neutralising antibodies that bind to the S2 subunit are less common and tend to be of lower potency than those that bind to Spike S1 (32, 33).

The main aim of this study was to investigate the phenotypic impact of naturally occurring Spike mutations in contemporary MERS-CoV strains. Naturally occurring mutations were identified by aligning MERS-CoV Spike sequences from human clinical isolates collected between 2012 – 2024 or from viruses that had been passaged in human cells. Fifteen mutations of interest were selected for further characterisation based on their location within the Spike protein, mutation frequency and inclusion in previous studies. The mutations selected included SNPs located in the NTD, RBD and adjacent to the S1/S2 cleavage site. Mutations were introduced to a wildtype Spike backbone, which was representative of the Spike proteins of clade B, lineage 5 viruses circulating in the Middle East. A lentiviral pseudotyping system was then used to investigate the impact of the naturally occurring mutations on entry and neutralisation (34, 35).

Improving our understanding of genotype to phenotype changes in MERS-CoV Spike is of particular importance due to the pandemic potential of this virus. Since emerging in 2012, frequent spillover events have occurred and several instances of human-to-human transmission have been identified (36, 37). The most recent examples are nine cases of MERS-CoV being reported to the World Health Organisation (WHO) in April 2025, with seven cases originating from one index case (38). Additionally, in early December 2025, two MERS-CoV cases were reported in travellers returning to France from the Arabian Peninsula (39). This highlights the importance of continued investigation of contemporary MERS-CoV strains, as with each spillover event the virus continues to evolve and adapt (37), which may lead to increased infectivity and transmission in humans.

## Methods

### MERS-CoV Spike sequence alignment

MERS-CoV Spike sequences used in this study were obtained from GenBank (NCBI) (n = 571), the Saudi Arabian Public Health Authority (n = 1) and from our own in-house sequencing of reverse genetics viruses passaged in human cells (n = 12) using an amplicon-based sequencing approach that has previously been described (40). In total, 584 Spike sequences isolated from humans were aligned using the Multiple Sequence Alignment with MUSCLE tool in the Unipro UGENE software. The spike reference sequence used for this alignment was generated from a clinical isolate collected in Saudi Arabia in 2019 (Supplementary Figure 1). Spike amino acid changes relative to the reference and their frequency were identified and recorded. 15 SNPs of interest occurring in the NTD, RBD and adjacent to the S1/S2 cleavage site were selected for further characterisation. Mutations of interest were selected based on their frequency and locations within the Spike protein, as well as according to prior knowledge available from previous studies in the literature.

### Cell maintenance

Human embryonic kidney 293T (HEK-293T) cells and baby hamster kidney 21 (BHK-21) cells were purchased from the American Type Culture Collection (ATCC). Cells were cultured and maintained in Dulbecco’s Modified Eagle’s Medium – high glucose (DMEM; Sigma-Aldrich) supplemented with 10% fetal bovine serum (FBS; Sigma-Aldrich), termed complete DMEM, at 37°C with 5% CO_2_. Cells were passaged every 3 – 4 days.

### Plasmids

Codon optimised MERS-CoV Spike contained in the pcDNA3.1(+) expression vector was designed and synthesised using GeneArt™ (Thermo Fisher). The Spike sequence was taken from a clinical isolate collected in Saudi Arabia in 2019 and was representative of the MERS-CoV viruses that were circulating in the Middle East (Supplementary Figure 1). The Kozak sequence was included immediately upstream and a C-terminal FLAG-tag was included downstream of the Spike sequence. The final 16 amino acids, containing an endoplasmic reticulum (ER)/Golgi retention motif and an endosomal recycling motif, were removed from the C-terminal cytoplasmic tail of Spike to increase pseudotyping efficiency (41). This plasmid will be referred to as the wildtype Spike plasmid. No glycoprotein (GP) pcDNA3.1(+) vector containing a FLAG-tag (Addgene) was used as an empty vector control and VSV-G contained within the pcDNA3.1(+) vector was used as the positive control as it is known to pseudotype efficiently. The lentiviral pseudotyping system consisted of two plasmids; p8.91 and pCSFLW. p8.91 is a second-generation packaging construct containing the HIV-1 core genes, *gag-pol*, under the control of a human CMV promoter (42). The pCSFLW transfer plasmid contained the firefly luciferase reporter flanked by HIV-1 regulatory long terminal repeats (LTRs) and a packaging signal (43). GFP1-7 and GFP8-11 were contained within the pcDNA3.1(+) expression vector and were used in the split GFP cell-cell fusion assays. Human DPP4 in a pCMV3-untagged expression vector was purchased from Sino Biological.

### Plasmid propagation

Plasmids were propagated by transforming into NEB^®^ 5-alpha competent *Escherichia coli* (*E. coli*) (high efficiency) cells (New England Biolabs; NEB) via heat shock as detailed in the manufacturer’s instructions. Bacteria were then spread onto appropriate selective LB agar plates and incubated overnight at 37°C to allow *E. coli* colony formation. Single colonies were picked from the LB agar plates and expanded in 250mL selective LB media at 37°C and 250rpm overnight. Plasmid DNA was then purified from the *E.coli* cultures using the QIAGEN plasmid maxi kit (Qiagen) as per manufacturer’s instructions. Plasmids were quantified using the Nanodrop One spectrophotometer (Thermo Scientific) and verified via restriction endonuclease digest and Sanger sequencing (Eurofins Genomics) prior to use in this study.

### Site-directed mutagenesis

The plasmid containing the wildtype Spike sequence was used as a template for site-directed mutagenesis to produce plasmid variants carrying the mutations of interest. SNPs were introduced using the QuikChange Lightning site-directed mutagenesis kit (Agilent) as detailed in the manufacturer’s instructions. 50ng of wildtype spike plasmid was used as the template in each reaction, along with the appropriate mutagenesis primer pair for each mutation (Supplementary Table 1). Mutagenesis reactions were digested with 2µL DpnI and 2µL of each reaction was transformed into *E.coli* for plasmid propagation as described above.

### Transfection

Transfections were performed using Lipofectamine 3000 (Thermo Fisher) as detailed in the manufacturer’s instructions. Plasmids were transfected into HEK-293T cells to produce pseudotypes or for the split GFP cell-cell fusion assay, while plasmids were transfected into BHK-21 cells for the split GFP cell-cell fusion assay only. HEK-293T cells were seeded into 6-well tissue culture treated plates (CytoOne, Starlab) coated with poly-L lysine (Sigma-Aldrich) as per the manufacturer’s instructions. BHK-21 cells were seeded into 6-well tissue culture treated plates (CytoOne, Starlab). Cells were seeded at an appropriate density 24h prior to transfection to produce cell monolayers that were 70 – 80% confluent at the point of transfection. Transfections were performed in Opti-MEM™ media (Thermo Fisher) using 7.5µL Lipofectamine 3000, 5µL P3000 reagent and the appropriate plasmid concentrations described previously (34, 44, 45). After incubating for 15 minutes at room temperature, transfection mixes were added to the cells and incubated overnight at 37°C with 5% CO_2_. At 18h post-transfection, transfection mixes were removed and replaced with 3mL complete DMEM. Cell plates were then incubated at 37°C with 5% CO_2_ for a further 24 – 48h.

### Generation of pseudotypes

Pseudotypes were generated via co-transfection of the p8.91 packaging plasmid, the pCSFLW transfer plasmid and the different variants of the MERS-CoV Spike plasmid using Lipofectamine 3000 (Thermo Fisher) as described above. The plasmid concentrations used in each transfection reaction were 0.6µg p8.91, 0.6µg pCSFLW and 0.5µg Spike, as previously described (34, 35). Pseudotypes were harvested at 48 and 72 hours post-transfection and pooled. Pseudotypes were centrifuged at 300g for 10 minutes at 4°C, before being aliquoted and stored at -80°C. Pseudotype cell lysates were also collected for use in Western blot analyses by adding 250µL Laemmli buffer (Sigma-Aldrich) combined with 250µL PBS (Sigma-Aldrich) to each well and pipetting up and down to disrupt the cell monolayer.

### Titration assays

Titration and viability of pseudotypes was tested using a luciferase-based infectivity assay as previously described (34). Four days prior to pseudotype infection, HEK-293T cells were seeded at an appropriate density into tissue culture treated 6 well plates (CytoOne, Starlab) coated with poly-L lysine (Sigma-Aldrich), as described above. After 24h, HEK-293T cells were transfected with 0.5µg human DPP4 using Lipofectamine 3000 (Thermo Fisher) as described above. At 18 hours post-transfection, media was replaced with complete phenol-free DMEM and cell plates were incubated for a further 24h to allow for DPP4 expression.

On the day prior to pseudotype infections, HEK-293T DPP4 cells were washed and collected by pipetting complete phenol-free DMEM up and down to disrupt the cell monolayer. Cells were counted and seeded into white tissue culture treated 96-well plates (Sigma-Aldrich) at a density of 2 x 10^4^ cells per well in complete phenol-free DMEM and incubated at 37°C with 5% CO_2_ for 24h. On the day of pseudotype infections, all pseudotypes were serially diluted 10-fold from neat – 1 x 10^-4^ in phenol-free DMEM – high glucose (Sigma-Aldrich) supplemented with 2% FBS (Sigma-Aldrich). 100µL of the serially diluted pseudotypes were added to each well and plates were incubated at 37°C with 5% CO_2_ for 72h. After 72h, media was removed and 50µL Bright-Glo™ (Promega) substrate combined 1:1 with phenol-free DMEM was added per well and incubated in the dark for five minutes at room temperature. Luciferase activity was then measured using a GloMax® Discover microplate reader (Promega) with a 0.5s integration time.

### Split GFP cell-cell fusion assays

Split GFP cell-cell fusion assays were used to investigate the entry properties of the Spike proteins carrying different mutations and were performed as previously described (34, 45). Three days prior to the split GFP cell-cell fusion assay, HEK-293T and BHK-21 cells were seeded at appropriate densities in tissue culture treated 6 well plates (CytoOne, Starlab), as described above. After 24h, HEK-293T cells were co-transfected with 0.25µg Spike and 0.5µg GFP1-7 and BHK-21 cells were co-transfected with 0.5µg human DPP4 and 0.5µg GFP8-11 using Lipofectamine 3000 (Thermo Fisher), as described above. After replacing the media with complete phenol-free DMEM 18h post-transfection, cell plates were incubated for a further 24h to allow for protein expression.

Split GFP cell-cell fusion assays were then performed by co-culturing HEK-293T cells expressing Spike and GFP1-7 with BHK-21 cells expressing the DPP4 receptor and GFP8-11. On the day of co-culturing, media was removed, cells were washed and collected by pipetting phenol-free DMEM (Sigma-Aldrich) up and down to disrupt the cell monolayer. Cells were counted and seeded into a 96-well plate (CytoOne, Starlab) at a density of 5 x 10^4^ cells/well in 100µL phenol-free DMEM – high glucose (Sigma-Aldrich) supplemented with 2% FBS (Sigma-Aldrich). Cells were incubated at 37°C with 5% CO_2_ for 1h to allow cells to settle and adhere to the plate. After 1h, cell plates were transferred to an IncuCyte® (Sartorius) where they were incubated at 37°C with 5% CO_2_ and GFP expression and syncytia formation were measured from images taken every 2h for 24h.

### Microneutralisation assays

Microneutralisation assays were used to investigate whether the Spike mutations impacted neutralisation by pooled patient sera and were performed as previously described (34, 44). Pooled patient sera used in these assays was the 1^st^ WHO International Standard for Anti-MERS-CoV Immunoglobulin G (Human) 19/178 purchased from the National Institute for Biological Standards and Control (NIBSC). It contained the pooled serum from two individuals from South Korea who had recovered from MERS-CoV infection (46).

Three days prior to pseudotype infection, HEK-293T cells were seeded at an appropriate density into tissue culture treated 6 well plates (CytoOne, Starlab) coated with poly-L lysine (Sigma-Aldrich), as described above. After 24h, HEK-293T cells were transfected with 0.5µg human DPP4 using Lipofectamine 3000 (Thermo Fisher) as described above. At 18 hours post-transfection, media was replaced with complete phenol-free DMEM and cell plates were incubated for a further 24h to allow for DPP4 expression. Pooled patient sera was then diluted 1:10 in phenol-free DMEM – high glucose (Sigma-Aldrich) supplemented with 2% FBS (Sigma-Aldrich) and serially diluted 2-fold to 1:2560 in white tissue culture treated 96-well plates (Sigma-Aldrich). Pseudotypes at a titre equivalent to 10^5^ – 10^6^ relative light units (RLU) were then combined 1:1 with the pooled patient sera and incubated for 1 hour at 37°C with 5% CO_2_. During this incubation, HEK-293T DPP4 cells were washed and collected in phenol-free DMEM – high glucose (Sigma-Aldrich) supplemented with 2% FBS (Sigma-Aldrich) by pipetting up and down. After incubating the pseudotypes and sera for 1 hour, cells were seeded at a density of 2 x 10^4^ cells per well in the white 96-well plates containing the pseudotypes and sera. Plates were then incubated for 72h at 37°C with 5% CO_2_. After 72h, media was removed and replaced with 50µL Bright-Glo™ (Promega) substrate combined 1:1 with phenol-free DMEM – high glucose (Sigma-Aldrich). Plates were incubated in the dark for five minutes at room temperature prior to luciferase activity being measured using a GloMax® Discover microplate reader (Promega) with a 0.5s integration time.

### Statistical analysis

Data were tested for normal and log-normal distribution to dictate the statistical tests used. Subsequent analyses are described in the figure legends. All graphs were generated in GraphPad Prism, version 10. Statistical analyses were performed in GraphPad Prism, version 10.

## Results

### Identification of mutations of interest in MERS-CoV Spike protein

Naturally occurring MERS-CoV Spike mutations identified in human infections may result in phenotypic differences in virus entry and neutralisation. Improved virus entry or partial immune escape could pose a significant risk to human health. To investigate this, naturally occurring amino acid mutations were identified by aligning the Spike genes from 584 human MERS-CoV sequences. A representative clade B, lineage 5 virus collected in Saudi Arabia in 2019 was used as the reference wildtype Spike sequence (Supplementary Figure 1). This clinical isolate sequence reflects the viruses currently circulating in the Middle East and contains 5 amino acid changes in its Spike sequence relative to the EMC/2012 reference strain.

Twenty seven SNPs were identified in the Spike genes in this alignment, and their frequencies were recorded (Figure 1A). The majority of the mutations identified were located in the RBD (24) (amino acids 381 – 588; Figure 1A – D). G94R, Q98R and Q304R were located in the NTD, while L745F and L746K were located adjacent to the S1/S2 cleavage site. L411F, F473S and I529T were the most commonly occurring mutations, with 42, 22 and 26 isolates containing each of these mutations, respectively (Figure 1A). Mutations of interest were then selected for further characterisation based on location, frequency and the likelihood of impacting entry and/or neutralisation properties of the virus. The following 15 mutations were down selected: G94R, Q98R, Q304R, T387P, L411F, T424I, F473S, L506F, D510G, I529T, E536K, W553R, T560I, L745F and T746K (Figure 1; Table 1). These mutations were found independently in separate isolations of the virus and never together. Therefore, in this study, mutations were characterised as single point mutations only.

**Figure 1:**
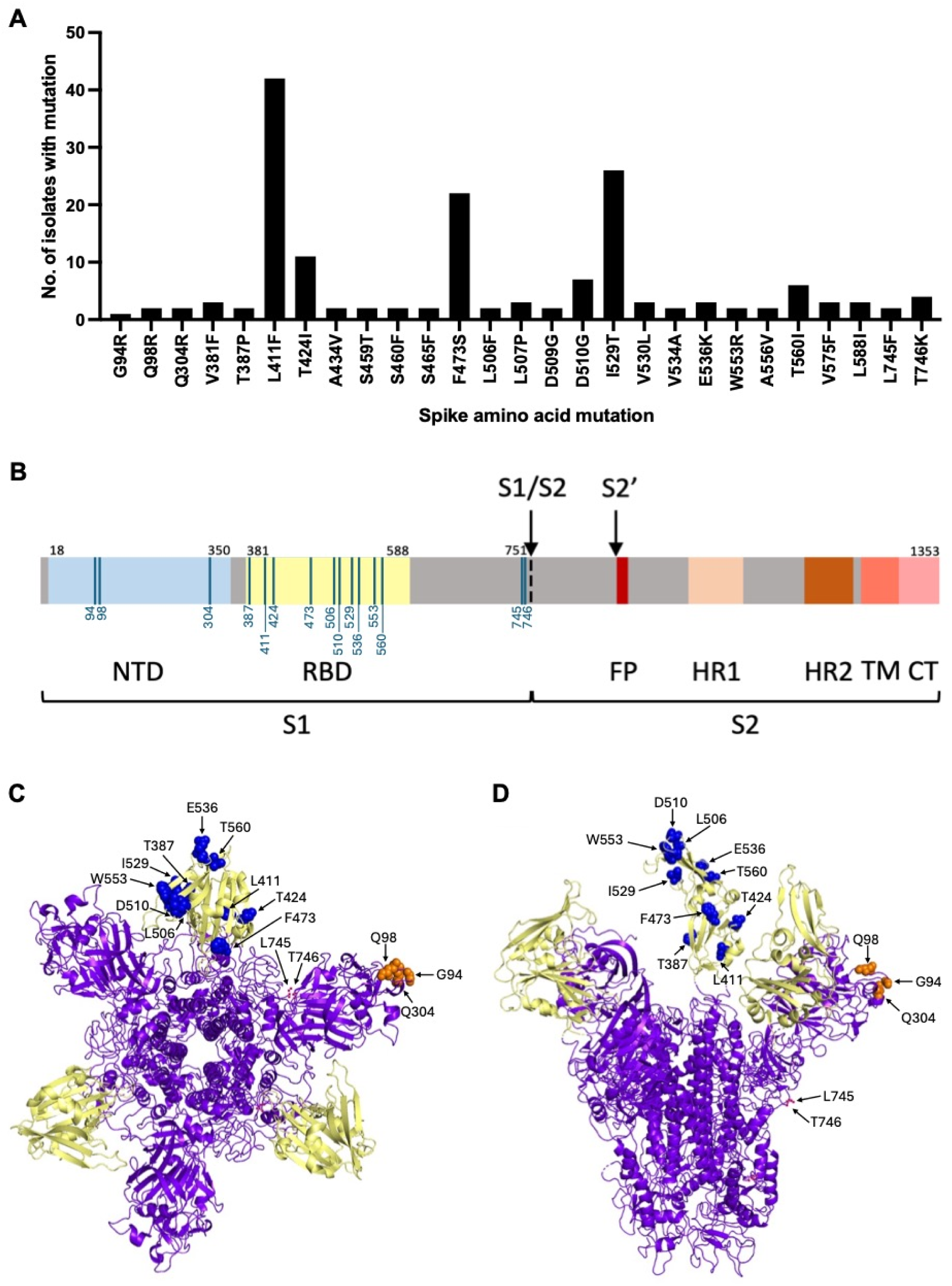
Summary of Spike mutations identified in an alignment of 584 MERS-CoV clinical isolates. (A) Mutations that were identified are shown on the x-axis and have been plotted against the number of clinical isolates carrying the mutation. (B) Schematic showing the different domains of Spike with the locations of the mutations that were selected for further characterisation labelled. Mutations were situated in the N-terminal domain (NTD; blue), receptor binding domain (RBD; yellow) and adjacent to the S1/S2 cleavage site (black arrow). S2’ cleavage site (S2’), fusion peptide (FP), heptad repeat region 1 (HR1), heptad repeat region 2 (HR2), transmembrane domain (TM) and cytoplasmic tail (CT) are also depicted. (C – D) Top and side views of the Spike protein structure in a pre-fusion state (Protein databank structure: 5X59; https://doi.org/10.2210/pdb5X59/pdb) with the locations of the mutations of interest highlighted. RBD is shown in yellow with the locations of the RBD mutations highlighted in blue. Locations of NTD mutations are highlighted in orange and cleavage site mutations are in pink.

**Table 1.** Summary of Spike mutations of interest that were selected for characterisation. Pseudotypes were assigned a mutant number and the location of each mutation is listed in the table below. The wildtype carried a representative clade B Spike sequence, which had five amino acid changes relative to the EMC/2012 reference strain. Mutants were generated via site-directed mutagenesis of the wildtype spike plasmid.

| <u>Pseudotype</u> | <u>Spike mutation</u> | <u>Location of mutation</u> |
| --- | --- | --- |
| Wildtype | Clinical sample<br>(Saudi Arabia, 2019) | 5 spike aa changes vs<br>EMC/2012 strain |
| Mutant 1<br>T387P | T387P | RBD |
| Mutant 2<br>L411F | L411F | RBD |
| Mutant 3<br>T424I | T424I | RBD |
| Mutant 4<br>F473S | F473S | RBD |
| Mutant 5<br>L506F | L506F | RBD |
| Mutant 6<br>D510G | D510G | RBD |
| Mutant 7<br>I529T | I529T | RBD |
| Mutant 8<br>E536K | E536K | RBD |
| Mutant 9<br>W553R | W553R | RBD |
| Mutant 10<br>T560I | T560I | RBD |
| Mutant 11<br>L745F | L745F | Adjacent to S1/S2 |
| Mutant 12<br>T746K | T746K | Adjacent to S1/S2 |
| Mutant 13<br>G94R | G94R | NTD |
| Mutant 14<br>Q98R | Q98R | NTD |
| Mutant 15<br>Q304R | Q304R | NTD |

The locations of the 15 mutations of interest were mapped to show their positions relative to the different Spike domains and locations within the Spike protein structure (Figure 1B – D). Mutations located in the RBD were likely to directly interact with the DPP4 receptor, and therefore impact entry. Additionally, many of the RBD residues are exposed on the outer surface of the Spike S1 subunit (Figure 1C and 1D). This suggested that the RBD residues could be targeted by neutralising antibodies and mutations occurring in these positions may therefore impact neutralisation. Mutations located in the NTD were of interest due to their possible role in host cell attachment and their exposed locations on the outer surface of the Spike S1 subunit (Figure 1C and 1D) (27–29), making them possible targets for neutralising antibodies. Mutations located close to the S1/S2 cleavage site were selected on the basis that they may alter cleavage efficiency and therefore impact entry. L745 and T746 were also located on the outer surface of the Spike protein (Figure 1D) and may therefore impact neutralisation if this region is targeted by antibodies.

### Generation of MERS-CoV Spike lentiviral pseudotypes

To evaluate the effect of different mutations on entry and neutralisation properties, MERS-CoV Spike lentiviral pseudotypes were generated by co-transfection of the lentiviral pseudotyping system and Spike expression plasmid into HEK-293T cells. This resulted in Spike being incorporated into nascent lentiviral particles to produce replication-deficient pseudotypes. Pseudotypes were then subsequently used to study entry and neutralisation properties of Spike proteins carrying the different mutations of interest.

The plasmids used to generate pseudotypes in this study were the p8.91 packaging plasmid, pCSFLW transfer plasmid and plasmids containing the Spike variants. Spike variant plasmids containing the mutations of interest were generated by site-directed mutagenesis of the wildtype Spike plasmid. VSV-G was included as a positive control for pseudotype generation and titration as it is known to pseudotype efficiently and no GP empty pcDNA3.1(+) vector (Addgene) was included as a negative control. Plasmids were verified by restriction endonuclease digest and submitted to Eurofins Genomics for Sanger sequencing, which confirmed that all Spike plasmid variants contained the expected mutations (data not shown). The lentiviral pseudotyping system and Spike plasmids were then used to generate pseudotypes via co-transfection into HEK-293T cells followed by harvesting of cell supernatants containing the pseudotypes after 48h and 72h.

After pseudotypes were harvested, cell lysates were collected for analysis by Western blot to confirm expression of Spike, the HIV-1 lentiviral *gag* and *pol* core genes, the firefly luciferase reporter and VSV-G. Western blot analysis indicated that protein expression was as expected for each of the pseudotype cell lysates (Supplementary Figure 2 – 4). Spike expression was confirmed in all pseudotype cell lysates, except for the VSV-G and no GP controls. Expression of VSV-G was only observed in the VSV-G pseudotype cell lysate. Expression of the HIV-1 core and the firefly luciferase reporter was observed for all pseudotypes, including VSV-G and no GP controls. GAPDH was included as a loading control and was detected across all samples including the no plasmid transfection control, as expected. Spike, the HIV-1 lentiviral core and the firefly luciferase reporter were not found to be expressed in the no plasmid transfection control cell lysate. This indicated that protein expression was as expected for all pseudotypes and controls.

### Titres of pseudotypes carrying mutations in the Spike NTD were reduced relative to the wildtype

Once correct protein expression had been confirmed, pseudotypes were titrated in a luciferase-based infectivity assay to verify that they were functional. Functional pseudotypes should be able to enter HEK-293T DPP4 cells via binding to the DPP4 receptor, which will subsequently result in expression of the firefly luciferase reporter. Spike wildtype, Spike mutants 1 – 15, VSV-G and no GP pseudotypes were serially diluted 10-fold and used to infect HEK-293T DPP4 cells. After 72h, luminescence was measured to determine pseudotype titres, with the no GP negative control being indicative of background (Figure 2A – C). Undiluted pseudotypes with titres >2.0 Log_10_ RLU above the no GP negative control were considered to be infective and therefore functional (34).

**Figure 2:**
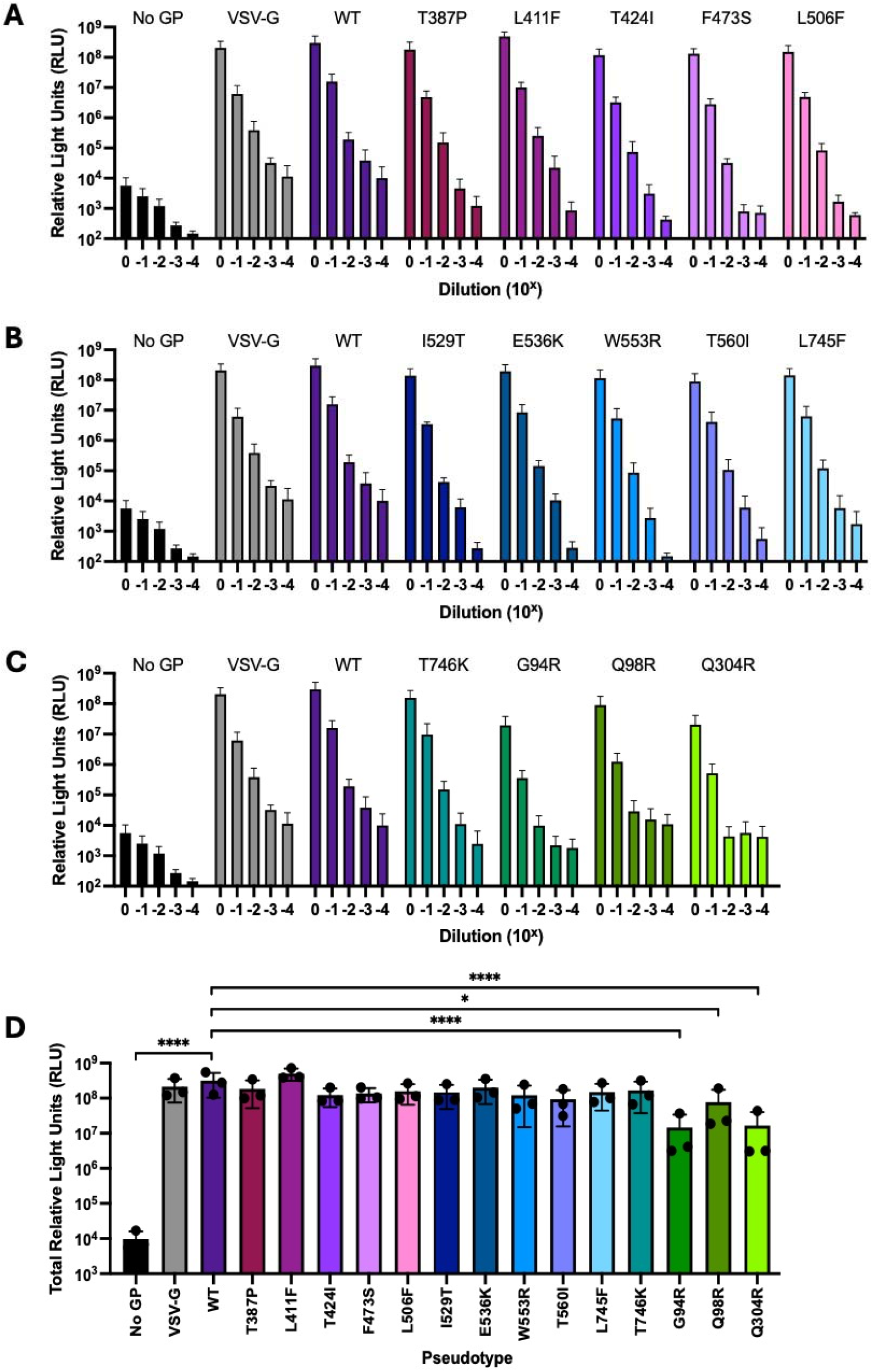
Titres of pseudotypes carrying NTD mutations were reduced relative to the wildtype. Pseudotypes were serially diluted 10-fold and used to infect HEK-293T cells expressing DPP4. (A – C) Luciferase activity was measured 72h post-infection using a GloMax® Discover microplate reader (Promega). (D) Total luciferase activity across the dilution series was plotted to include all dilutions simultaneously in the statistical analysis. One-way ANOVA with Dunnet’s multiple comparisons test indicated that titres of no GP, G94R, Q98R and Q304R were significantly reduced relative to the wildtype (n=3 independent replicates; error bars show mean with standard deviation; *p<0.05, ****p<0.0001).

Pseudotype titres were then expressed as total RLU to generate a single data point per pseudotype in each experiment for statistical analysis, while still using all data points across the dilution series (Figure 2D). Total RLU titres for mutants 1 – 15 and the positive and negative controls were compared to the wildtype Spike pseudotype. Pseudotypes carrying the NTD mutations G94R, Q98R or Q304R were found to have significantly reduced titres relative to the wildtype (p<0.0001, p<0.05 and p<0.0001, respectively). The titres of the remaining pseudotypes (mutants 1 – 12) were comparable to the wildtype. The titre of the no GP negative control was significantly reduced relative to the wildtype Spike pseudotype (p<0.0001), as expected.

### I529T, E536K and L745F improved entry relative to the wildtype

Split GFP cell-cell fusion assays were performed to assess the impact of the Spike mutations on receptor binding and membrane fusion. The NTD has been shown to play a role in host cell attachment (27–29), meaning that the mutations located in the NTD (G94R, Q98R and Q304R) may impact entry. Mutations located in the RBD (T387P, L411F, T424I, F473S, L506F, D510G, I529T, E536K, W553R and T560I) may alter receptor binding properties. Mutations located near the S1/S2 cleavage site (L745F and T746K) may impact cleavage efficiency, which would have a knock-on effect on virus entry.

To set up the split GFP cell-cell fusion assays, HEK-293T cells expressing wildtype Spike or a Spike variant and GFP1-7 were co-cultured with BHK-21 cells expressing the DPP4 receptor and GFP8-11 (Figure 3A). Spike binds to the DPP4 receptor, subsequently resulting in membrane fusion and syncytia formation. The two halves of GFP can then be assembled into a functional protein, emitting fluorescent signal corresponding to the receptor binding and membrane fusion ability of the Spike protein. GFP expression was measured from images taken of the co-cultured cells every 2h for 24h using an IncuCyte® (Sartorius) (Figure 3B – C). HEK-293T cells expressing VSV-G (glycoprotein negative control) and no GP pcDNA3.1(+) (no glycoprotein negative control) were also included in this experiment. Background fluorescence measurements taken from HEK293T cells expressing Spike and GFP1-7 only and BHK-21 cells expressing DPP4 and GFP8-11 only (not co-cultured) were averaged and subtracted from the values for the co-cultured conditions.

**Figure 3:**
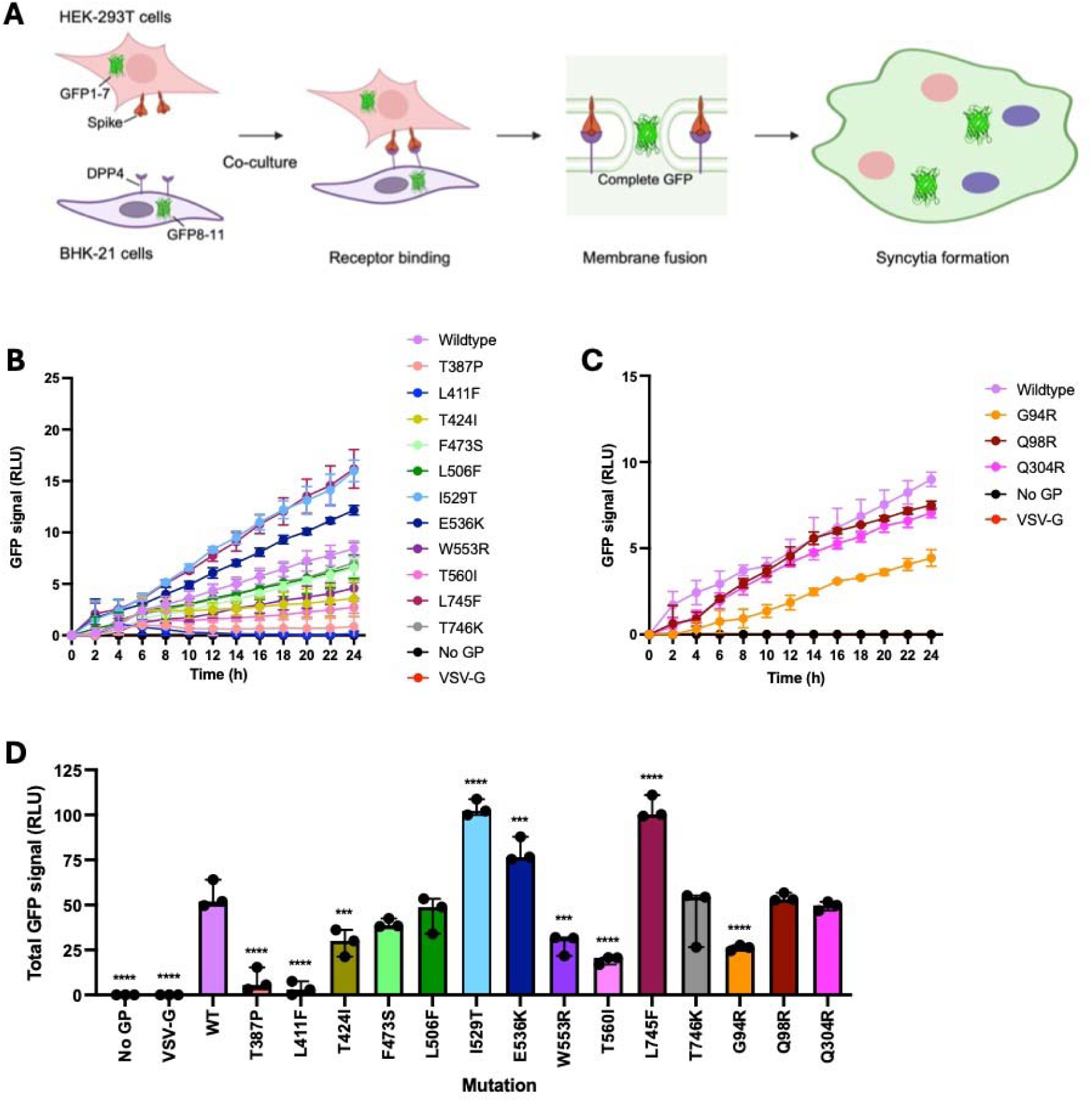
Receptor binding and membrane fusion of I529T, E536K and L745F mutants were improved relative to the wildtype. (A) Schematic depicting the formation of syncytia during the split GFP cell-cell fusion assay. HEK-293T cells expressing Spike and GFP1-7 were co-cultured with BHK-21 cells expressing DPP4 and GFP8-11. Spike binds the DPP4 receptor leading to membrane fusion and syncytia formation, resulting in the two halves of GFP being assembled into a functional fluorescent protein. (B – C) GFP signal emitted by the Spike mutants and controls over the 24h period following co-culturing, normalised to background controls. GFP signal was measured every 2h using an IncuCyte® (Sartorius). (D) Total GFP signal across the time course was plotted to include all time points simultaneously in the statistical analysis. One-way ANOVA with Dunnet’s multiple comparisons test indicated that I529T, E536K and L745F significantly improved receptor binding and membrane fusion relative to the wildtype. T387P, L411F, T424I, W553R, T560I and G94R significantly reduced receptor binding and membrane fusion relative to the wildtype (n=3 independent replicates; error bars show mean with standard deviation; ***p<0.001, ****p<0.0001).

Fluorescence readings were then expressed as total RLU to generate a single data point per Spike protein in each experiment for statistical analysis, while still using all data points from the time course (Figure 3D). Total RLU titres for mutants 1 – 15 and the controls were compared to the wildtype Spike protein. Entry properties were significantly improved for the Spike proteins carrying I529T (p<0.0001), E536K (p<0.001) and L745F (p<0.0001) when compared to the wildtype. In contrast, entry properties were significantly reduced for the Spike proteins carrying T387P (p<0.0001), L411F (p<0.0001), T424I (p<0.001), W553R (p<0.001), T560I (p<0.0001) and G94R (p<0.0001) relative to the wildtype. Entry properties of the Spike proteins carrying F473S, L506F, T746K, Q98R and Q304R were comparable to the wildtype. No receptor binding and membrane fusion was observed for the VSV-G and no GP negative controls, as expected.

Syncytia formation was also visualised using the images taken on the IncuCyte® (Sartorius) and normalised to the whole plate. Representative fluorescence images taken at 0h, 6h, 12h, 18h and 24h are shown for each experimental condition (Figure 4 and 5). Syncytia formation for each Spike protein increased over time, indicating that receptor binding and membrane fusion was increasing over the 24h experiment. Spike proteins carrying I529T, E536K and L745F demonstrated increased syncytia formation and GFP expression due to improved entry properties (Figure 5). Spike proteins carrying T387P, L411F, T424I, W553R, T560I and G94R demonstrated decreased syncytia formation and GFP expression relative to the wildtype due to reduced receptor binding and membrane fusion (Figure 4 and 5).

**Figure 4:**
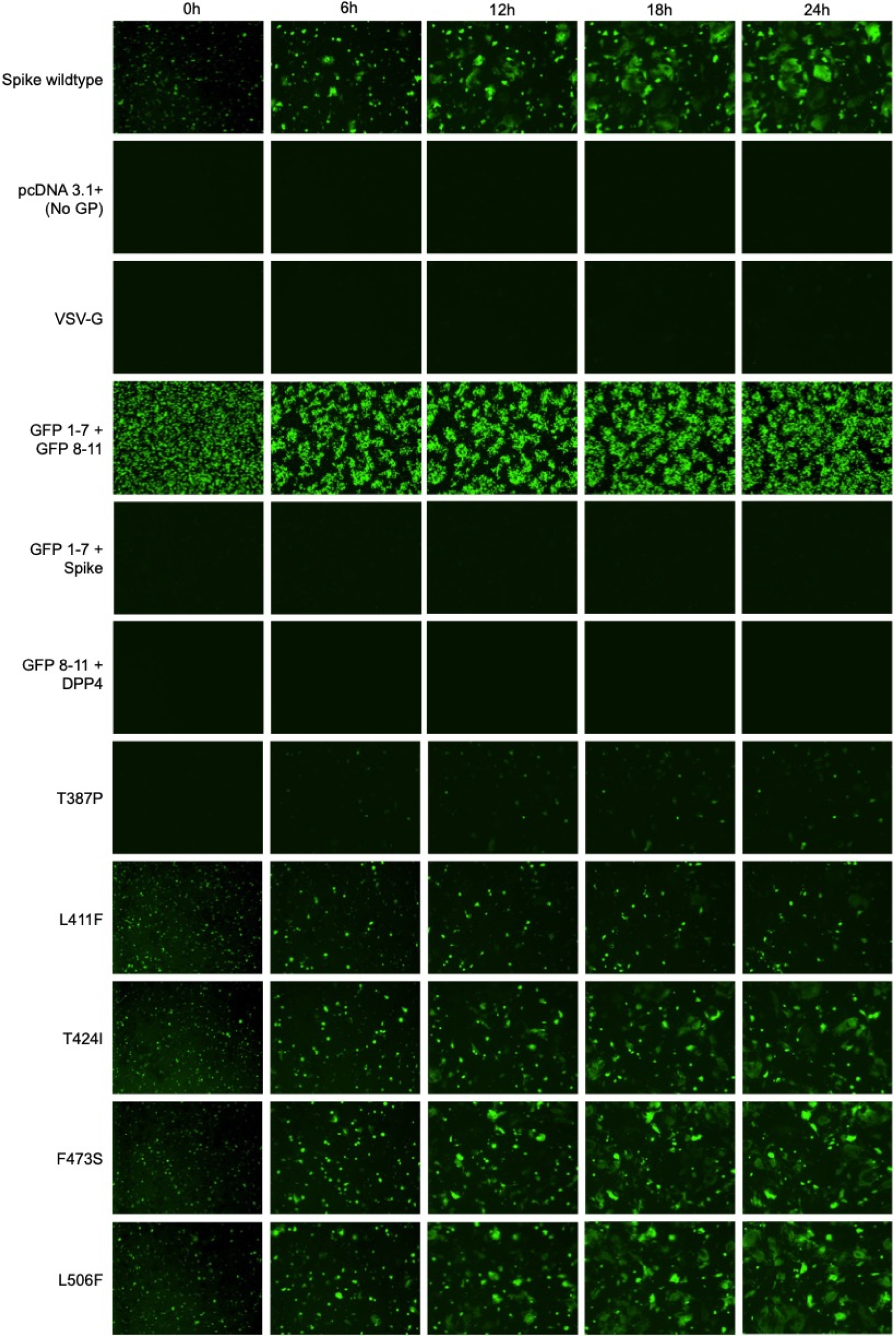
T387P, L411F and T424I mutants demonstrated decreased syncytia formation when compared to the wildtype. HEK-293T cells expressing GFP1-7 and either wildtype Spike, Spike mutant 1 – 5, pcDNA3.1(+) (no GP) or VSV-G were co-cultured with BHK-21 cells expressing the DPP4 receptor and GFP8-11. GFP1-7 + GFP8-11, GFP1-7 + Spike and GFP8-11 + DPP4 were included as controls and to measure background signal for normalisation. GFP signal was measured from images taken every 2h on the IncuCyte® (Sartorius). Representative images taken at 0h, 6h, 12h, 18h and 24h post-co-culture, with GFP signal normalised to the whole plate, are shown here.

**Figure 5:**
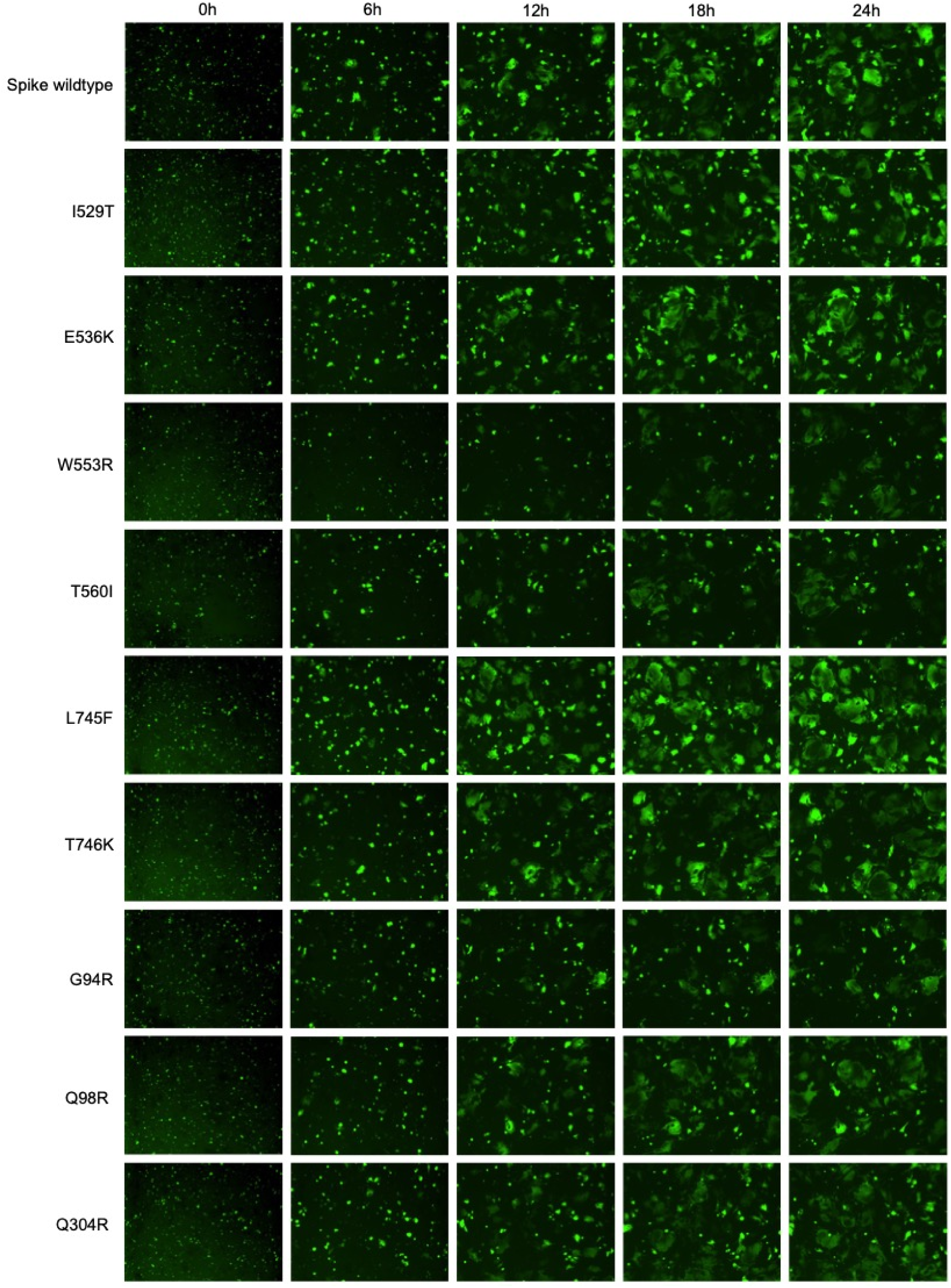
I529T, E536K and L745F mutants demonstrated increased syncytia formation when compared to the wildtype. HEK-293T cells expressing GFP1-7 and either wildtype Spike or Spike mutant 7 – 15 were co-cultured with BHK-21 cells expressing the DPP4 receptor and GFP8-11. GFP signal was measured from images taken every 2h on the IncuCyte® (Sartorius). Representative images taken at 0h, 6h, 12h, 18h and 24h post-co-culture, with GFP signal normalised to the whole plate, are shown here.

Importantly, no syncytia formation or GFP expression was observed for the VSV-G and pcDNA3.1(+) no GP controls (Figure 4). GFP1-7 + Spike only and GFP8-11 + DPP4 only controls were included for background fluorescent measurements and no syncytia formation or GFP expression were observed in the images for these conditions. As expected, GFP expression, but not syncytia formation, was observed for GFP1-7 + GFP8-11 co-transfected into HEK-293T cells as a GFP expression control (Figure 4).

### Pseudotypes carrying L411F, T424I, L506F, L745F and T746K were neutralised less efficiently by pooled patient sera than the wildtype

Microneutralisation assays were performed to assess the impact of the Spike mutations on neutralisation by pooled patient sera. The NTD and RBD are known to contain epitopes recognised by neutralising antibodies (15, 47, 48), meaning that mutations located in these regions may contribute to partial immune escape. Mutations located close to the S1/S2 site may impact cleavage efficiency, which could alter which epitopes are exposed (49). Alternatively, these mutations may be part of antibody epitopes that are not yet understood.

To set up the microneutralisation assays, pooled patient sera from recovered patients that had been infected with MERS-CoV (NIBSC, (46)) was serially diluted 2-fold from 1:20 – 1:5120. Spike wildtype, Spike mutants 1 – 15, VSV-G and no GP pseudotypes were added at a titre of 10^5^ – 10^6^ RLU and incubated for 1h with the sera. No sera controls were also included for each pseudotype. HEK-293T DPP4 cells were then added and incubated for a further 72h. Antibodies in the pooled patient sera should bind to the Spike proteins of the pseudotypes, preventing their entry into HEK-293T DPP4 cells and resulting in reduced expression of the firefly luciferase reporter. After 72h, luminescence was measured to determine the extent by which the pseudotypes were inhibited by the pooled patient sera (Figure 6).

**Figure 6:**
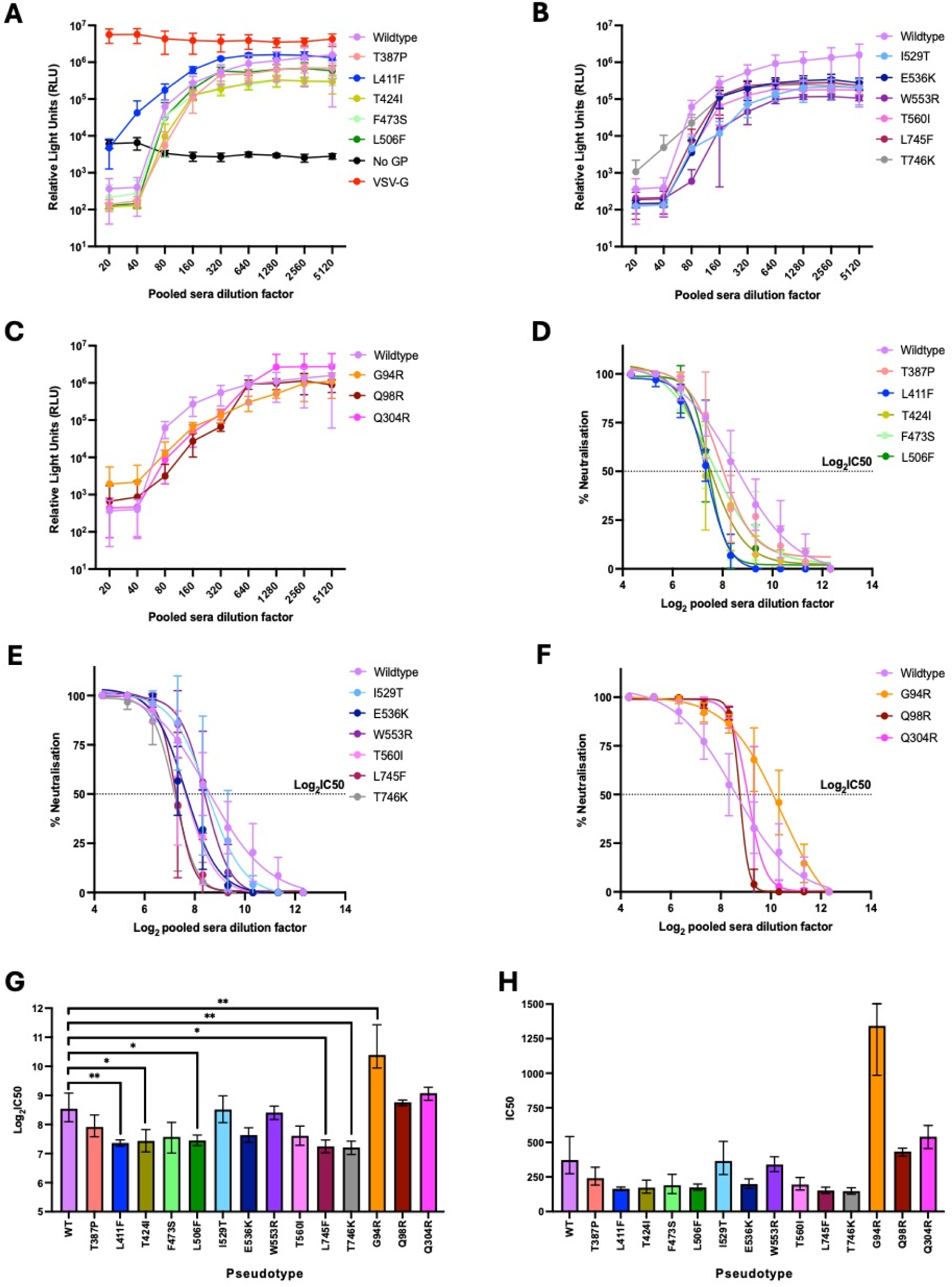
Pseudotypes carrying L411F, T424I, L506F, L745F and T746K mutations were neutralised less efficiently than the wildtype. Pooled patient sera was serially diluted two-fold from 1:20 – 1:5120. Spike pseudotypes were diluted to 10^5^ – 10^6^ RLU based on titres determined previously. MERS-CoV Spike pseudotypes, no GP and VSV-G pseudotype controls were incubated with pooled patient sera for 1h prior to addition of HEK-293T cells expressing DPP4. (A – C) After 72h, luciferase activity was measured using a GloMax® Discover microplate reader (Promega) and raw relative light units (RLU) data were plotted against dilution factor for each pseudotype. (D – F) % Neutralisation was calculated from the raw data using no sera control (0% value) and mean no GP control (100% value). Log(inhibitor) vs normalized response – variable slope non-linear regression curves were fitted to the % neutralisation values. (G) Log_2_IC_50_ values were calculated from the log(inhibitor) vs response – variable slope curves. One-way ANOVA with Dunnet’s multiple comparisons test indicated that pseudotypes carrying L411F, T424I, L506F, L745F and T746K mutations were neutralised less efficiently than wildtype, while G94R was neutralised more efficiently relative to the wildtype (n=4 independent replicates, error bars show mean values with standard deviation; *p<0.05, **p<0.01). (H) Log_2_IC_50_ values were anti-logged to give the half maximal inhibitory dilution factor of pooled patient sera for each pseudotype.

Percent neutralisation values were calculated from the raw data using the no sera controls for each pseudotype as the 0% neutralisation RLU value (Figure 6A – F). The mean no GP control RLU value was set as the 100% neutralisation value. Non-linear regression curves were fitted to the percent neutralisation values using GraphPad Prism v10 software (Figure 6D – F). Log_2_ 50% inhibitory concentration (Log_2_IC_50_) values were then calculated from the non-linear regression curves (Figure 6G), as previously described (50). Log_2_IC_50_ values were then anti-logged to give the IC_50_ values (Figure 6H). IC_50_ values represent pooled patient sera dilution factor that results in 50% inhibition of each pseudotype.

Statistical analysis of Log_2_IC_50_ values was performed to compare neutralisation of mutants 1 – 15 with the wildtype Spike pseudotype. Pseudotypes carrying L411F, T424I, L506F, L745F and T746K mutations were neutralised less efficiently by pooled patient sera than the wildtype (p<0.01 T424I, T746K; p<0.05 T424I, L506F, L745F) (Figure 6G). The pseudotype carrying G94R was neutralised more efficiently than the wildtype pseudotype (p<0.01). Microneutralisation of the remaining Spike mutant pseudotypes was comparable to the wildtype. Importantly, pooled patient sera had no inhibitory effect on VSV-G and no GP controls (Figure 6A).

## Discussion

This study was conducted to investigate the phenotypic impact of naturally occurring Spike mutations in contemporary MERS-CoV strains. 15 naturally occurring Spike mutations were selected for characterisation using a contemporary MERS-CoV Spike backbone. The mutations characterised were SNPs located in the NTD, RBD or adjacent to the S1/S2 cleavage site. Novel naturally occurring mutations impacting entry and neutralisation of contemporary MERS-CoV were identified in this study. I529T, E536K and L745F mutations improved MERS-CoV entry (Figure 3 and 5). Whereas L411F, T424I, L506F, L745F and T746K were shown to increase resistance to neutralisation by pooled patient sera (Figure 6).

Prior to this study, knowledge surrounding the 15 mutations was limited due to having not previously been characterised, or characterised using an EMC/2012 backbone. Kleine-Weber et al previously demonstrated that L411F and F473S did not impact entry when introduced to the EMC/2012 Spike sequence in a VSV-based pseudotyping system (23). Additionally, L411F was shown not to impact virus neutralisation by sera isolated from patients that had been infected with MERS-CoV in 2014 (7). Spike residues 506, 536 and 553 have been confirmed to make direct contact with DPP4 during receptor binding (51), indicating that mutations occurring at these residues may impact entry. Tang et al demonstrated that the L506F substitution caused a minimal reduction in DPP4 binding and was neutralised less efficiently by antibodies isolated from a human antibody phage display library (52). However, it is important to note that EMC/2012 sequence was used as the Spike backbone and to generate the neutralising antibodies used in this study. Residues 506, 536 and 553 were also identified as being part of the epitope recognised by the MERS-GD27 neutralising antibody isolated from a recovered patient (47). Furthermore, when the L506F and E536K mutations were introduced to the EMC/2012 Spike sequence, neutralisation efficiency of MERS-GD27 was significantly decreased. Previous studies have also shown that I529T, one of the mutations associated with the MERS-CoV outbreak in South Korea in 2015, was associated with reduced affinity for the DPP4 receptor, reduced neutralisation with patient sera and reduced pathogenicity in mice (23, 53–55). Hoffman et al demonstrated that introducing the T746K polymorphism into the EMC/2012 Spike sequence in a VSV-based pseudotyping system resulted in increased S1/S2 cleavage, without increasing entry into Calu3 cells (56). However, when the E32A and Q1020R mutations, which are present in clade B viruses, were introduced to the EMC/2012 Spike sequence alongside T746K, entry into Calu3 cells was improved. This highlights the importance of relevant Spike backbone sequences when characterising contemporary virus strains due to mutations acting in a compensatory manner.

Luciferase-based titration assays showed that G94R, Q98R and Q304R mutations significantly reduced pseudotype titres relative to the wildtype (Figure 2D). This could indicate impaired entry of these mutants, as these mutations are located in the NTD, which is known to be involved in host cell attachment (27–29). However, it is also possible that discrepancies in transfection efficiency and/or packaging of Spike during pseudoparticle generation may have contributed to the differences in titres. Split GFP cell-cell fusion assays were therefore performed to further characterise the entry properties of the Spike variants.

Receptor binding and membrane fusion were enhanced for Spike proteins carrying I529T, E536K and L745F mutations (Figure 3). This was also reflected by increased syncytia formation for these Spike variants relative to the wildtype (Figure 5). I529T was previously shown to diminish entry (53), although the reduction in entry caused by this mutation was later demonstrated to only occur in cells with low levels of DPP4 expression (23). It is likely that the BHK-21 DPP4 cells used in this experiment expressed enough DPP4 for receptor binding and membrane fusion to occur. Given that residue 536 makes direct contact with the DPP4 receptor (51), the improved receptor binding and membrane fusion observed for the E536K mutant in this assay may be due to improved DPP4 binding. The L745F mutation is located in close proximity to the S1/S2 cleavage site, suggesting that the improvement in receptor binding and membrane fusion observed here may be due to altered S1/S2 cleavage efficiency of this Spike variant. More efficient cleavage at S1/S2 results in more Spike proteins on the pseudoparticle surface being primed for receptor binding, which has previously been demonstrated to increase host cell entry (26).

Receptor binding and membrane fusion were reduced for Spike variants carrying T387P, L411F, T424I, W553R, T560I and G94R mutations (Figure 3). This was also reflected by the reduced syncytia formation observed for these Spike variants relative to the wildtype (Figure 4 and 5). Spike residues 387, 411 and 424 are not known to directly interact with the DPP4 receptor. However, these substitutions may have caused minor changes in the Spike protein structure due to changes in size and charge of the amino acid side chains, resulting in a knock-on effect on receptor binding. In contrast to the reduction in receptor binding and membrane fusion observed for the L411F variant in this study, Kleine-Weber et al previously showed that L411F did not impact entry into HEK-293T DPP4 cells (23). This discrepancy may be attributed to the differences in Spike backbone used in each of these studies. The L411F variant used here includes five additional Spike amino acid changes associated with clade B, lineage 5 viruses, relative to the EMC/2012 Spike sequence that was used in the study conducted by Kleine-Weber et al.

Residues 553 and 560 are known to make direct contact with DPP4, which indicates that the reduced receptor binding and membrane fusion observed for the W553R and T560I variants may be due to reduced affinity for the receptor. In addition to reduced receptor binding/membrane fusion, the G94R variant also had a significantly lower titre than the wildtype (Figure 2 and 3). Given that residue 94 is located on the outer surface of the Spike NTD (Figure 1) and that G94R is associated with a change in amino acid charge, it is possible that the G94R mutation interferes with host cell attachment, resulting in the impaired entry properties observed for this Spike variant.

Pseudotypes carrying the L411F, T424I, L506F, L745F and T746K SNPs were neutralised less efficiently by pooled patient sera relative to the wildtype (Figure 6). In contrast to the results presented here, Schroeder et al previously showed that L411F did not impact virus neutralisation with patient sera (7). This difference may be due to the patient sera being obtained from patients infected with different clade B lineages. Schroeder et al used patient sera collected in 2014, prior to the emergence of lineage 5 viruses, whereas the pooled patient sera used in this study was obtained from patients infected with lineage 5 viruses during the South Korean MERS-CoV outbreak in 2015. This is also likely to be the reason differences in neutralisation for the I529T variant were not observed in this study, which does not align with previous studies (23, 54). The I529T mutation arose during the MERS-CoV outbreak in South Korea and was associated with reduced neutralisation relative to sera raised against EMC/2012 and early clade B viruses. However, the pooled patient sera used in this study was obtained from patients infected with lineage 5 strains that were likely carrying the I529T substitution.

T424I was neutralised less efficiently by pooled patient sera relative to the wildtype (Figure 6). This residue has not previously been identified as being located within an antibody epitope, although its location on the outer surface of the RBD (Figure 1) indicates that it could be targeted by neutralising antibodies. The L506F and E536K mutations have previously been shown to reduce neutralisation by sera obtained from patients infected with MERS-CoV during the South Korean outbreak in 2015 (47). These residues have been confirmed to be located within neutralising antibody epitopes and E536K was found to reduce neutralisation to a lesser extent than L506F (47, 52). In this study, L506F significantly reduced neutralisation by pooled patient sera (Figure 6), restoring the ability of this mutant to bind DPP4 and infect host cells. E536K also appeared to reduce neutralisation, although this was found not to be statistically significant in this study (Figure 6).

L745F and T746K pseudotype variants were neutralised less efficiently by pooled patient sera relative to the wildtype (Figure 6). Residues 745 and 746 are located within close proximity to the S1/S2 cleavage site (Figure 1), and S1/S2 cleavage can occur during biosynthesis of the Spike protein, or upon interaction with target host cells (25, 26). Differences in cleavage efficiency will impact the proportion of primed versus unprimed Spike proteins in the pseudotype variants, which may alter the epitopes exposed to neutralising antibodies (49). Interestingly, L745F was also found to increase receptor binding and membrane fusion in this study, which means that this substitution appears to have made MERS-CoV more infectious and provided increased resistance to neutralisation (Figure 3 and 6).

To summarise, a contemporary MERS-CoV pseudotyping system was developed and the phenotypic consequences of naturally occurring Spike mutations on receptor binding, membrane fusion and neutralisation were identified. This pseudotyping system can continue be used to characterise Spike mutations as they arise or could be used in the development of medical countermeasures for MERS-CoV. Continued investigation of genotype to phenotype consequences in MERS-CoV is essential for pandemic preparedness and reducing the risk to public health.

## Acknowledgements

We are grateful to Mohammed Alsayer (Public Health Authority, Riyadh, Saudi Arabia) for providing the wildtype MERS-CoV Spike sequence used in this study.

## Author contributions

Conceptualization: RD, WA and JAH.

Methodology: RD, HG, JN, NT and DB.

Validation: RD, HG, JN, NT and DB.

Formal analysis: RD, HG.

Investigation: RD, HG, JN, NT, DB, JPS, WA and JAH.

Resources: JN, NT and DB.

Data curation: RD.

Writing – original draft: RD and JAH.

Writing - Review and Editing: RD, NT, TMac, TM, RO, DB, WA, JAH.

Visualization: RD.

Supervision: DB, JPS, WA and JAH.

Project Administration: TMac, TM, RO and JAH.

Funding acquisition: DB, JPS, WA and JAH.

