## Supplementary Info for "Characterisation of naturally occurring MERS-CoV Spike mutations and their impact on entry and neutralisation"

**Supplementary Methods**

**Western blot**

Proteins were transferred onto a polyvinylidene difluoride (PVDF) membrane using Trans-Blot® Turbo™ mini PVDF transfer packs (BioRad) and the Trans-Blot® Turbo™ transfer system (BioRad) using the manufacturer’s settings for 1 Mini-PROTEAN® TGX gel per tray. Membranes were then blocked in 5% non-fat milk powder dissolved in tris-buffered saline with tween-20 (TBS-T; 20mM Tris, 150mM of NaCl, 0.1% Tween 20, pH 7.6) for 1h at room temperature. Membranes were incubated with appropriate primary antibodies diluted in blocking buffer at 4°C overnight (Supplementary Table 2). Prior to incubation with secondary antibodies, membranes were washed three times with TBS-T for 5 minutes per wash. Appropriate horseradish peroxidase conjugated secondary antibodies were diluted in blocking buffer and incubated with membranes for 1h at room temperature (Supplementary Table 2). Membranes were washed five times with TBS-T for 5 minutes per wash prior to the addition of enhanced chemiluminescent (ECL) substrate (BioRad) and imaging with a ChemiDoc imaging system (BioRad). After initial probing, membranes were stripped at room temperature as follows: two 10 minute incubations with stripping buffer (15g Glycine, 1g SDS, 10mL Tween 20, made up to 1L with double-distilled water d(dH_2_0); pH 2.2), two 10 minute washes in PBS (Sigma-Aldrich) and two five minute washes in TBS-T. Membranes were then blocked and probed for the GAPDH loading control as described above.

### **Supplementary Tables and Figures**

***Supplementary Table 1: Spike mutagenesis primer sequences.*** *Mutations were introduced to the wildtype spike plasmid via site-directed mutagenesis using the primers listed in the table.*

| **Pseudotype** | **Primer** | **Primer Sequence 5’ – 3’** |
| --- | --- | --- |
| Mutant 1  T387P | T387P_F  T387P_R | AATGCGACTTCAGCCCCCTGCTGTCTGGC  GCCAGACAGCAGGGGGCTGAAGTCGCATT |
| Mutant 2  L411F | L411F_F  L411F_R | GTTCACCAACTGCAATTACAACTTTACCAAGCTGCTGAGCC  GGCTCAGCAGCTTGGTAAAGTTGTAATTGCAGTTGGTGAAC |
| Mutant 3  T424I | T424I_F  T424I_R | TCCGTGAACGACTTCATCTGTAGCCAGATCAGC  GCTGATCTGGCTACAGATGAAGTCGTTCACGGA |
| Mutant 4  F473S | F473S_F  F473S_R | AACTACAAGCAGTCCTCCAGCAACCCTACCTGC  GCAGGTAGGGTTGCTGGAGGACTGCTTGTAGTT |
| Mutant 5  L506F | L506F_F  L506F_R | ATCAACAAGTGCAGCAGATTTCTGAGCGACGACAGAACC  GGTTCTGTCGTCGCTCAGAAATCTGCTGCACTTGTTGAT |
| Mutant 6  D510G | D510G_F  D510G_R | ACTGCTGAGCGACGGCAGAACCGAAGTGC  GCACTTCGGTTCTGCCGTCGCTCAGCAGT |
| Mutant 7  I529T | I529T_F  I529T_R | CCCTTGCGTGTCCACCGTGCCTAGCAC  GTGCTAGGCACGGTGGACACGCAAGGG |
| Mutant 8  E536K | E536K_F  E536K_R | CTAGCACCGTTTGGAAGGACGGCGACTAC  GTAGTCGCCGTCCTTCCAAACGGTGCTAG |
| Mutant 9  W553R | W553R_F  W553R_R | GGAAGGCGGAGGACGGCTGGTGGC  GCCACCAGCCGTCCTCCGCCTTCC |
| Mutant 10  T560I | T560I_F  T560I_R | GCTGGTGGCCTCTGGATCTATAGTGGCCATGA  TCATGGCCACTATAGATCCAGAGGCCACCAGC |
| Mutant 11  L745F | L745F_F  L745F_R | GCTCTGCCAGATACACCCATCACATTTACCCCAAGATCCG  CGGATCTTGGGGTAAATGTGATGGGTGTATCTGGCAGAGC |
| Mutant 12  T746K | T746K_F  T746K_R | GCCAGATACACCCATCACACTGAAACCAAGATCCGTG  CACGGATCTTGGTTTCAGTGTGATGGGTGTATCTGGC |
| Mutant 13  G94R | G94R_F  G94R_R | CGGACACGCCACCCGGACCACACCTCAGA  TCTGAGGTGTGGTCCGGGTGGCGTGTCCG |
| Mutant 14  Q98R | Q98R_F  Q98R_R | CGGCACCACACCTCGGAAACTGTTCGTGG  CCACGAACAGTTTCCGAGGTGTGGTGCCG |
| Mutant 15  Q304R | Q304R_F  Q304R_R | GCATCCGGTCCATCCGGAGCGACAGAAAAGC  GCTTTTCTGTCGCTCCGGATGGACCGGATGC |

**Supplementary Table 2: Primary and secondary (*) antibodies used for Western blot analysis.** Antibodies targeting the different pseudotype proteins were used at the dilutions listed in the table to confirm expression.

| **Target** | **Vendor** | **Cat. number** | **Dilution** |
| --- | --- | --- | --- |
| MERS-CoV Spike S1 | Thermo Fisher | PA5-119581 | 1/10000 |
| Firefly luciferase | Abcam | Ab185923 | 1/2000 |
| HIV-1 p24 | Abcam | Ab32352 | 1/2000 |
| VSV-G | GenScript | A00199 | 1/4000 |
| GAPDH | Abcam | Ab8245 | 1/5000 |
| Goat anti-rabbit* | Sigma | A6154 | 1/5000 |
| Goat anti-mouse* | Sigma | A4416 | 1/5000 |

***Supplementary Figure 1:* *Wildtype Spike sequence of representative MERS-CoV clade B virus isolated from a patient in Saudi Arabia in 2019.***

ATGATACACTCAGTGTTTCTACTGATGTTCTTGTTAACACCTACAGAAAGTTACGTTGATGTAGGGCCAGATTCTGTTAAGTCTGCTTGTATTGAGGTTGATATACAACAGACTTTCTTTGATAAAACTTGGCCTAGGCCAATTGATGTTTCTAAGGCTGACGGTATTATATACCCTCAAGGCCGTACATATTCTAACATAACTATCACTTATCAAGGTCTTTTTCCCTATCAGGGAGACCATGGTGATATGTATGTCTACTCTGCAGGACATGCTACAGGCACAACTCCACAAAAGTTGTTTGTAGCTAACTATTCTCAGGACGTCAAACAGTTTGCTAATGGGTTTGTCGTCCGTATAGGAGCAGCTGCCAATTCCACTGGCACTGTTATTATTAGCCCATCTACCAGCGCTACTATACGAAAAATTTACCCTGCTTTTATGCTGGGTTCTTCAGTTGGTAATTTCTCAGATGGTAAAATGGGCCGCTTCTTCAATCATACTCTAGTTCTTTTGCCCGATGGATGTGGCACTTTACTTAGAGCTTTTTATTGTATTCTAGAGCCTCGCTCTGGAAATCATTGTCCTGCTGGCAATTCCTATACTTCTTTTGCCACTTATCACACTCCTGCAACAGATTGTTCTGATGGCAATTACAATCGTAATGCCAGTCTGAACTCTTTTAAGGAGTATTTTAATTTACGTAACTGCACCTTTATGTACACTTATAACATTACCGAAGATGAGATTTTAGAGTGGTTTGGCATTACACAAACTGCTCAAGGTGTTCACCTCTTCTCATCTCGGTATGTTGATTTGTACGGCGGCAATATGTTTCAATTTGCCACCTTGCCTGTTTATGATACTATTAAGTATTATTCTATCATTCCTCACAGTATTCGTTCTATCCAAAGTGATAGAAAAGCTTGGGCTGCCTTCTACGTATATAAACTTCAACCGTTAACTTTCCTGTTGGATTTTTCTGTTGATGGTTATATACGCAGAGCTATAGACTGTGGTTTTAATGATTTGTCACAACTCCACTGCTCATATGAATCCTTCGATGTTGAATCTGGAGTTTATTCAGTTTCGTCTTTCGAAGCAAAACCTTCTGGCTCAGTTGTGGAACAGGCTGAAGGTGTTGAATGTGATTTTTCAACTCTTCTGTCTGGCACACCTCCTCAGGTTTATAATTTCAAGCGTTTGGTTTTTACCAATTGCAATTATAATCTTACCAAATTGCTTTCACTTTTTTCTGTGAATGATTTTACTTGTAGTCAAATATCCCCAGCAGCAATTGCTAGCAACTGTTATTCTTCACTGATTTTGGATTATTTTTCATACCCACTTAGTATGAAATCCGATCTCAGTGTTAGTTCTGCTGGTCCAATATCCCAGTTTAATTATAAACAGTCTTTTTCTAATCCCACTTGTTTGATTTTAGCGACTGTTCCTCATAACCTTACTACTATTACTAAGCCTCTTAAGTACAGCTATATTAACAAGTGCTCTCGTCTTCTTTCTGATGATCGTACTGAAGTACCTCAGTTAGTGAACGCTAATCAATACTCACCCTGTGTATCCATTGTCCCATCCACTGTGTGGGAAGACGGTGATTATTATAGGAAACAACTATCTCCACTTGAAGGTGGTGGCTGGCTTGTTGCTAGTGGCTCAACTGTTGCCATGACTGAGCAATTACAGATGGGCTTTGGTATTACAGTTCAATATGGTACAGACACCAATAGTGTTTGCCCCAAGCTTGAATTTGCTAATGACACAAAAATTGCCTCTCAATTAGGCAATTGCGTGGAATATTCCCTCTATGGTGTTTCGGGCCGTGGTGTTTTTCAGAATTGCACAGCTGTAGGTGTTCGACAGCAGCGCTTTGTTTATGATGCGTACCAGAATTTAGTTGGCTATTATTCTGATGATGGCAACTACTACTGTTTGCGTGCTTGTGTTAGTGTTCCTGTTTCTGTCATCTATGATAAAGAAACTAAAACCCACGCTACTCTATTTGGTAGTGTTGCATGTGAACACATTTCCTCTACCATGTCTCAATACTCCCGTTCTACGCGATCAATGCTTAAACGGCGAGATTCTACATATGGTCCCCTTCAGACACCTGTTGGTTGTGTCCTAGGAATTGTTAATTCCTCTTTGTTCGTAGAGGACTGCAAGTTGCCTCTTGGTCAATCTCTCTGTGCTCTTCCTGACACACCTAGTACTCTCACACCTCGCAGTGTGCGCTCTGTTCCAGGTGAAATGCGCTTGGCATCCATTGCTTTTAATCATCCTATTCAGGTTGATCAACTTAATAGTAGTTATTTTAAATTAAGTATACCTACTAATTTTTCCTTTGGTGTGACTCAGGAGTACATTCAGACAACCATTCAGAAAGTTACTGTTGATTGTAAACAGTACGTTTGCAATGGTTTCCAGAAGTGTGAGCAATTACTGCGCGAGTATGGCCAGTTTTGTTCCAAAATAAACCAGGCTCTCCATGGTGCCAATTTACGCCAGGATGATTCTGTACGTAATTTGTTTGCGAGCGTGAAAAGCTCTCAATCATCTCCTATCATACCAGGTTTTGGAGGTGACTTTAATTTGACACTTCTAGAACCTGTTTCTATATCTACTGGCAGTCGTAGTGCACGTAGTGCTATTGAGGATTTGCTATTTGACAAAGTCACTATAGCTGATCCTGGTTATATGCAAGGTTACGATGATTGTATGCAGCAAGGTCCAGCATCAGCTCGTGATCTTATTTGTGCTCAATATGTGGCTGGTTATAAAGTATTACCTCCTCTTATGGATGTTAATATGGAAGCCGCGTACACTTCATCTTTGCTTGGCAGCATAGCAGGTGTTGGCTGGACTGCTGGCTTATCCTCCTTTGCTGCTATTCCATTTGCACAGAGTATTTTTTATAGGTTAAACGGTGTTGGCATTACTCAACAGGTTCTTTCAGAGAACCAAAAGCTTATTGCCAATAAGTTTAATCAGGCTCTGGGAGCTATGCAAACAGGCTTCACTACAACTAATGAAGCTTTTCGGAAGGTTCAGGATGCTGTGAACAACAATGCACAGGCTCTATCCAAATTAGCTAGCGAGCTATCTAATACTTTTGGTGCTATTTCCGCCTCTATTGGAGACATCATACAACGTCTTGATGTTCTCGAACAGGACGCCCAAATAGACAGACTTATTAATGGCCGTTTGACAACACTAAATGCTTTTGTTGCACAGCAGCTTGTTCGTTCCGAATCAGCTGCTCTTTCCGCTCAATTGGCTAAAGATAAAGTCAATGAGTGTGTCAAGGCACAATCCAAGCGTTCTGGATTTTGCGGTCAAGGCACACATATAGTGTCCTTTGTTGTAAATGCCCCTAATGGCCTTTATTTTATGCATGTTGGTTATTACCCTAGCAACCACATTGAGGTTGTTTCTGCTTATGGTCTTTGCGATGCAGCTAACCCTACTAATTGTATAGCCCCTGTTAATGGCTACTTTATTAAAACTAATAACACTATGATTGTTGATGATTGGTCATATACTGGCTCGTCCTTCTATGCACCTGAGCCCATCACCTCTCTTAATACTAAGTATGTTGCACCACAGGTGACATACCAAAACATTTCTACTAACCTCCCTCCTCCTCTTCTCGGCAATTCCACCGGGATTGACTTCCAAGATGAGTTGGATGAGTTTTTCAAAAATGTTAGCACCAGTATACCTAATTTTGGTTCTCTAACACAGATTAATACTACATTACTCGATCTTACCTACGAGATGTTGTCTCTTCAACAAGTTGTTAAAGCCCTTAATGAGTCTTATATAGACCTTAAAGAGCTTGGCAATTATACTTATTACAACAAATGGCCGTGGTACATTTGGCTTGGTTTCATTGCTGGGCTTGTTGCCTTAGCTCTATGCGTCTTCTTCATACTGTGCTGCACTGGTTGTGGCACAAACTGTATGGGAAAACTTAAGTGTAATCGTTGTTGTGATAGATACGAGGAATACGACCTCGAGCCGCATAAGGTTCATGTTCACTAA

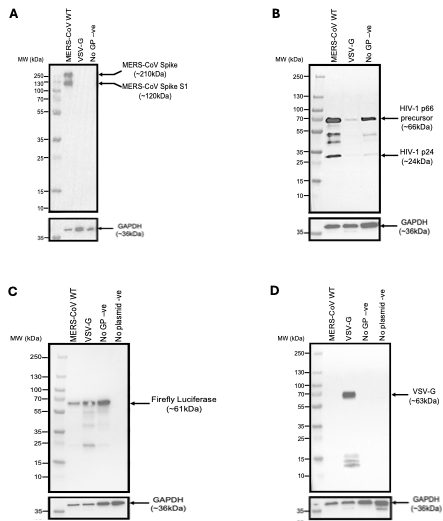

**Supplementary Figure 2: Western blot analysis of wildtype Spike pseudotype cell lysate confirmed expression of Spike protein, HIV-1 gag/pol lentiviral core proteins and the firefly luciferase reporter.** Cell lysates were harvested after generation of wildtype Spike, VSV-G positive control and no GP negative control pseudotypes. Cell lysates from a no plasmid transfection control were also harvested. Cell lysates were used for Western blot analysis to confirm expression of (A) Spike, (B) HIV-1 gag/pol lentiviral core, (C) firefly luciferase reporter and (D) VSV-G. GAPDH was also included as a loading control (A-D).

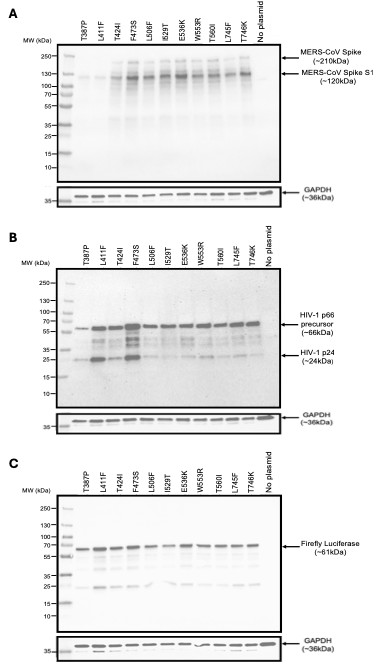

***Supplementary Figure 3: Western blot analysis of Spike pseudotype mutant 1 – 12 cell lysates confirmed expression of Spike protein, HIV-1 gag/pol lentiviral core proteins and the firefly luciferase reporter.*** *Cell lysates were harvested after generation of Spike pseudotype mutants 1 – 12. Cell lysates from a no plasmid transfection control were also harvested to be included as a negative control. Cell lysates were used for Western blot analysis to confirm expression of (A) Spike, (B) HIV-1 gag/pol lentiviral core and (C) the firefly luciferase reporter. (A-D) GAPDH was included as a loading control.*

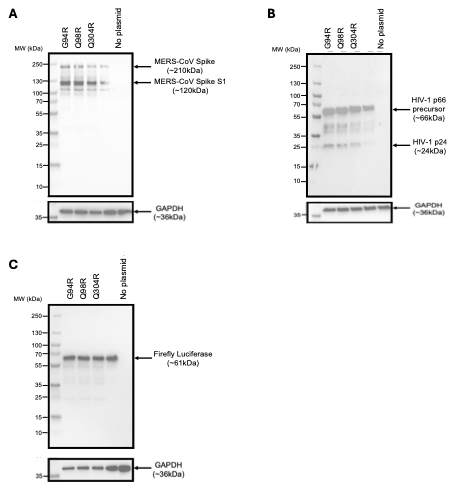

***Supplementary Figure 4: Western blot analysis of Spike pseudotype mutant 13 – 15 cell lysates confirmed expression of Spike protein, HIV-1 gag/pol lentiviral core proteins and the firefly luciferase reporter.*** *Cell lysates were harvested after generation of Spike pseudotype mutants 13 – 15. Cell lysates from a no plasmid transfection control were also harvested to be included as a negative control. Cell lysates were used in Western blot analysis to confirm expression of (A) Spike, (B) HIV-1 gag/pol lentiviral core and (C) the firefly luciferase reporter. GAPDH was also included as a loading control (A-D). The unlabelled samples on these blots are not relevant to this study.*
